# Single-cell RNA sequencing reveals microenvironment context-specific routes for epithelial-mesenchymal transition in pancreas cancer cells

**DOI:** 10.1101/2023.05.30.542969

**Authors:** Brooke A. Brown, Matthew J. Lazzara

## Abstract

In the PDAC tumor microenvironment, multiple factors initiate the epithelial-mesenchymal transition (EMT) that occurs heterogeneously among transformed ductal cells, but it is unclear if different drivers promote EMT through common or distinct signaling pathways. Here, we use single-cell RNA sequencing (scRNA-seq) to identify the transcriptional basis for EMT in pancreas cancer cells in response to hypoxia or EMT-inducing growth factors. Using clustering and gene set enrichment analysis, we find EMT gene expression patterns that are unique to the hypoxia or growth factor conditions or that are common between them. Among the inferences from the analysis, we find that the FAT1 cell adhesion protein is enriched in epithelial cells and suppresses EMT. Further, the receptor tyrosine kinase AXL is preferentially expressed in hypoxic mesenchymal cells in a manner correlating with YAP nuclear localization, which is suppressed by FAT1 expression. AXL inhibition prevents EMT in response to hypoxia but not growth factors. Relationships between FAT1 or AXL expression with EMT were confirmed through analysis of patient tumor scRNA-seq data. Further exploration of inferences from this unique dataset will reveal additional microenvironment context-specific signaling pathways for EMT that may represent novel drug targets for PDAC combination therapies.

## INTRODUCTION

In pancreatic ductal adenocarcinoma (PDAC), an especially aggressive tumor subtype exists that is enriched in mesenchymal characteristics. The mesenchymal phenotype is associated with decreased survival and increased resistance to therapy relative to other subtypes (1–4) and correlates with gene signatures for ductal cell epithelial-mesenchymal transition (EMT) (3). EMT occurs in response to varied initiating factors in the PDAC tumor microenvironment, including receptor ligands such as transforming growth factor beta (TGFβ), hepatocyte growth factor (HGF), and epidermal growth factor (5–7), and the low oxygen tension (hypoxia) resulting from PDAC tumor hypovascularity (3,8–10). It is unclear if different modes of EMT induction use distinct or common signaling pathways.

EMT occurs heterogeneously in PDAC tumors and cell populations (11–13). Even in cell culture conditions, where spatial variation in extrinsic EMT-driving factors is minimal, phenotypic heterogeneity is observed, and the degree of heterogeneity depends on cell background and mode of EMT induction (3). Heterogeneity is also affected by the durability of the transition, which is greater in response to hypoxia than growth factors (3). EMT heterogeneity endows differential chemoresistance to a subset of cells within tumors (12) and, as with any form of cell-to-cell heterogeneity, creates challenges for understanding the signaling or transcriptional processes that govern the phenotype (14). Prior work provides clues about the basis for EMT heterogeneity in different cancer cell settings, although generally without investigating fundamentally different classes of EMT-inducing conditions in parallel. In a colon cancer cell line subjected to single-cell RNA sequencing (scRNA-seq), variability in Wnt pathway activity due to intrinsic heterogeneity in the expression of key signaling network nodes correlates with the baseline mesenchymal state (15). Further, ATAC-seq revealed enrichment of the transcription factor RUNX2 in Wnt-high cells, which was confirmed via *in vivo* overexpression of RUNX2 promoting metastasis and in clinical samples, RUNX2 correlated to poor survival (15). In MCF10A breast cancer cells treated with TGFβ, scRNA-seq demonstrates that some cells activate different EMT signaling pathways in series while others activate them in parallel (with Notch playing a particularly important role), leading to variability among cells in the rate of EMT progression and contributing to apparent phenotypic heterogeneity in a snapshot view (16). In lung, prostate, breast, and ovarian cancer cells treated with a diverse set of growth factors including TGFβ, EGF, and TNF⍺, scRNA-seq reveals minimal overlap of differentially expressed genes despite some common changes in the expression of genes associated with EMT (e.g., *EPCAM, VIM*) or specific EMT-regulating signaling pathways (e.g., *JUN, EGFR*), suggesting distinct transcriptional and signaling regulatory programs even among ligands for receptor kinases (17).

Here, we performed scRNA-seq on a baseline epithelial pancreas cancer cell line that was treated with a combination of TGFβ and HGF or cultured under hypoxic conditions. Both conditions produced a heterogeneous EMT among cells, enabling the identification of regulatory transcriptional processes that lead to or restrain EMT in a context-specific manner using clustering and gene set enrichment analysis. Among the inferences from the analysis, we found that expression of the FAT1 cell adhesion protein and Hippo pathway activity, which is regulated by FAT1 (18) and suppresses YAP nuclear localization (19), are generally enriched in epithelial cells. We further identified the receptor tyrosine kinase AXL, which is transcriptionally regulated by YAP (20) and can drive EMT in other cancers (21,22), as a hypoxia-specific driver of EMT in pancreas cancer cells. This and other inferences from our analysis were confirmed in an analysis of PDAC patient tumors. In addition to these experimentally validated predictions, the dataset gathered here provides a wealth of additional inferences that can be pursued to understand the context-specific basis for EMT induction in a pancreas cancer cell background.

## RESULTS

### EMT occurs heterogeneously in response to growth factors or hypoxia

HPAF-II human pancreas cancer cells were driven to undergo EMT by treatment with exogeneous TGFβ+HGF or culture in 1% O_2_. The combination of growth factors for receptors that independently promote EMT was used to activate downstream effectors broadly and thereby increase the odds that regulatory processes unique to growth factors or hypoxia are identified. HPAF-II cells were used because they are baseline epithelial and an established model for well-differentiated PDAC (23). As we have observed previously (3), the extent of mesenchymal transition differed between growth factors and hypoxia as EMT drivers, with hypoxia promoting vimentin expression in only a minority of cells and growth factors causing most cells to express abundant vimentin (**Figure 1A**). Thus, both conditions produced a heterogeneous EMT among cells but to different degrees. These observations motivate the possibility that different molecular pathways are responsible for EMT in response to the different drivers and prompt exploration of the basis for heterogeneity of EMT for each condition.

**Figure 1.**
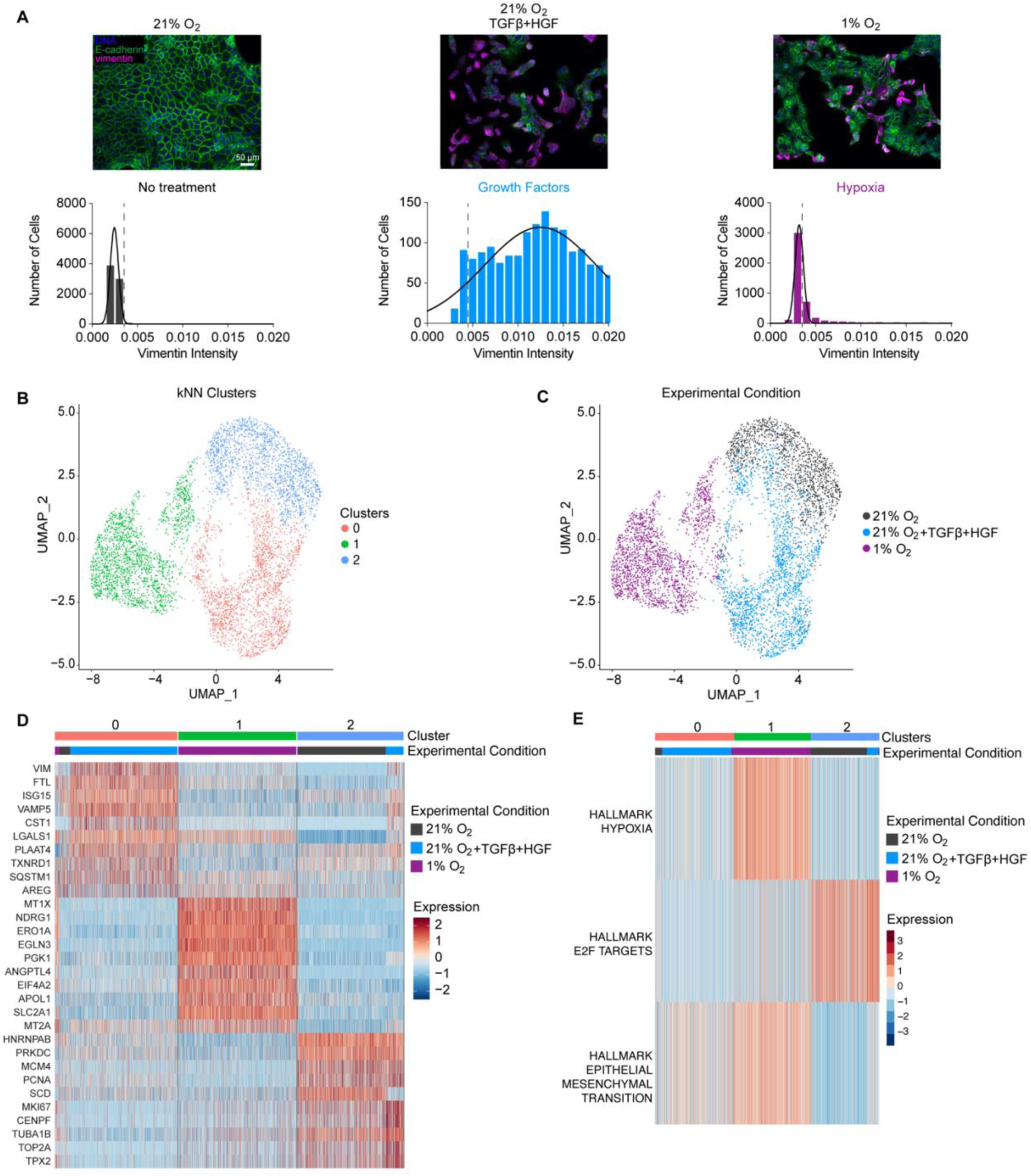
EMT is heterogenous across and within each treatment condition. **(A)** HPAF-II cells were cultured in 21% O_2_ with or without 10 ng/mL TGFβ and 50 ng/mL HGF or cultured in 1% O_2_ for 120 hr. Immunofluorescence microscopy was performed for the indicated proteins. Histograms show vimentin intensities across all biological replicates. *n* = 3. The dotted vertical line denotes the threshold for the intensity vimentin positivity. **(B)** UMAP projection of k-Nearest Neighbor (kNN) clustering performed on all genes for the aggregated scRNA-seq data from HPAF-II cells as treated in (A). **(C)** For the same UMAP projection shown in (B), cells are color-coded based on experimental condition. **(D)** The top 10 differentially expressed genes per cluster with respect to the other two clusters were calculated. **(E)** Gene set enrichment analysis was performed per cell for the 50 Hallmark gene sets. Displayed are the top three significantly differentially enriched gene sets among the three clusters as calculated by ANOVA.

Given that EMT is highly regulated by transcription factors (e.g., Slug, ZEB1) and the feasibility of single-cell transcriptomics, we utilized single-cell RNA-sequencing (scRNA-seq) to probe the transcriptional basis for EMT heterogeneity. We performed scRNA-seq on HPAF-II cells cultured in 21% O_2_ with or without TGFβ+HGF or cultured in 1% O_2_. As expected, cells cultured in 1% O_2_ were enriched in hypoxia-inducible factor (HIF) target genes (24) **(Supp Figure S1A,B)**, confirming that a hypoxic response occurred. Given the inherent technical noise in expression for any single gene in the data set, we performed principal component analysis (PCA) to allow for a group of genes to be used for subsequent clustering. The number of principal components (PC) for PCA projection was calculated by computing the variance in the dataset explained as a function of PC and using 0.1% as a cutoff **(Supp Figure S2A)**. Using a projection into 20 PCs, the aggregated data for all three conditions was clustered using k-nearest neighbors (kNN). The optimal number of clusters was determined by computing silhouette scores per cell for up to 10 clusters. Three clusters provided the highest mean silhouette score **(Supp Figure S2B)**. Clustered cells were then visualized in a UMAP projection (**Figure 1B**), and clusters corresponded well with treatment conditions (**Figure 1C**). To investigate the basis for cluster separation, we identified the top 10 differentially expressed genes (DEGs) by cluster (**Figure 1D**). Of note, DEGs for cluster 0 included the mesenchymal-associated genes *VIM* and *LGALS1*. For Cluster 1, DEGs of note included the hypoxia-associated genes *PGK1* and *SLC2A1*. For Cluster 2, DEGs of note included *MKI67* and *TUBA1B*, which play important roles in proliferation (a process most characteristic of the epithelial state). To identify the top differentially regulated cell processes among clusters, enrichment scores were calculated on a per cell basis for the 50 Hallmark Gene Sets (**Figure 1E**). The top three gene sets that were significantly differentially enriched among the three clusters include Hallmark Hypoxia (most expressed in Cluster 1), Hallmark E2F Targets (mostly expressed in Cluster 2), and Hallmark EMT (most expressed in Clusters 0 and 1). Thus, in addition to hypoxic response and proliferation, EMT represents one of the most significant overall differences among clusters, despite inter- and intra-condition differences in EMT induction.

### Differential gene expression between epithelial and mesenchymal cells reveals the governing signaling pathways

Given the relevance of EMT gene regulation in explaining variance among conditions, we next identified cells directly as epithelial or mesenchymal to probe the transcriptional differences between them. The full data set was re-clustered using the previously described approach but using features from a patient-derived pan-cancer EMT (pcEMT) signature (25), which contains 77 genes associated with epithelial or mesenchymal cell states and specifying two clusters (**Figure 2A**). This created one cluster enriched in epithelial (E) genes and one enriched in mesenchymal (M) genes (**Figure 2B**). Cells with no treatment (21% O_2_) overlapped primarily with the E cluster, while TGFβ+HGF-treated cells overlapped primarily with the M cluster; cells cultured at 1% O_2_ were roughly evenly split between the E and M clusters (**Figure 2C**). Differences between the clusters are also apparent in UMAP projections color-coded by the expression of the EMT markers *CDH1* (E-cadherin) and *VIM* (vimentin) (**Figure 2D**). As expected, the top 10 differentially expressed genes for the two clusters included cell-adhesion markers (i.e., *CDH17*, *CEACAM5*) and mesenchymal genes (i.e., *VIM* and *LGALS1*) (**Figure 2E**).

**Figure 2.**
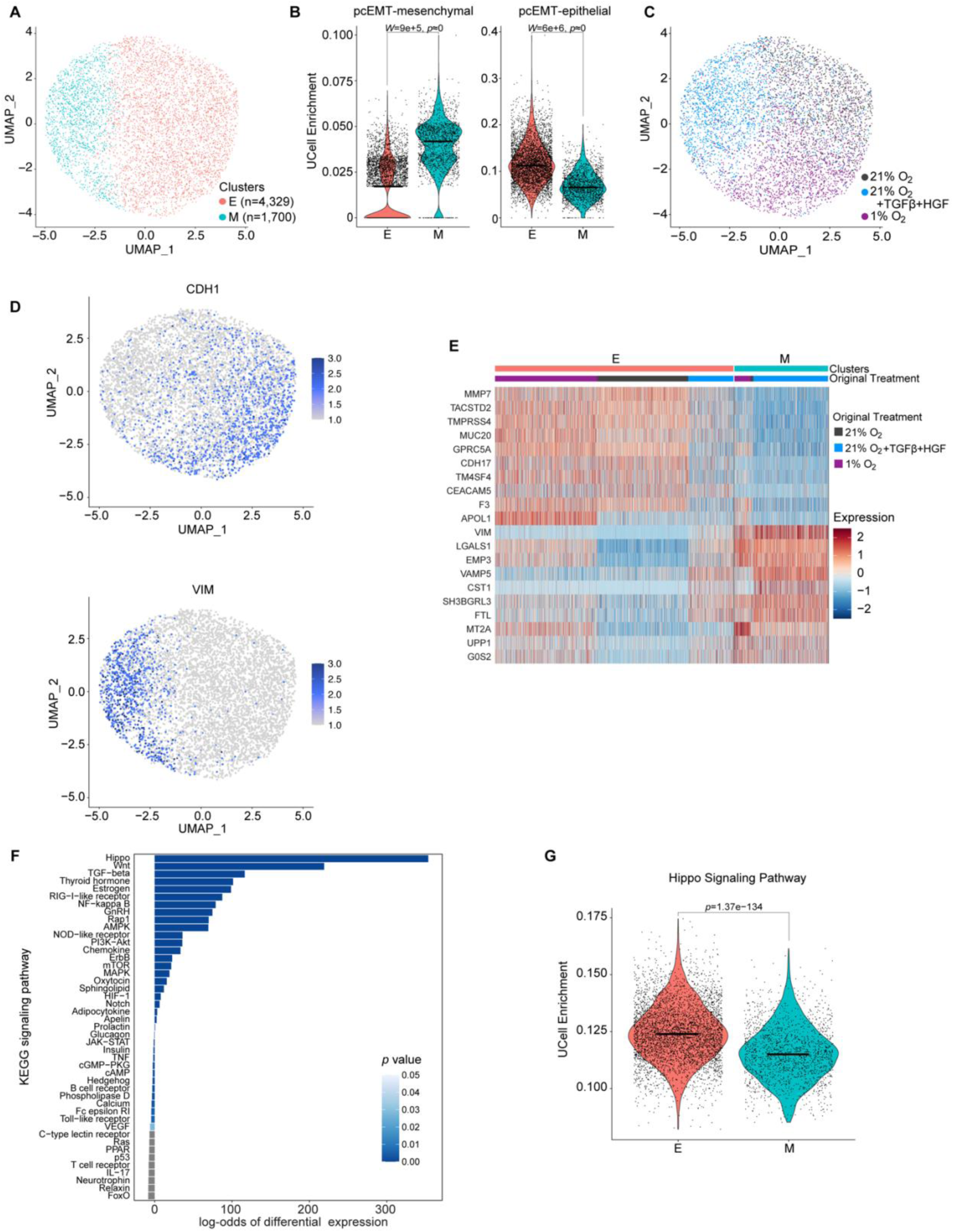
Cells were clustered on EMT-gene signature into epithelial and mesenchymal clusters to determine that the Hippo pathway is enriched in epithelial cells. **(A)** Aggregated scRNA-seq data from HPAF-II cells cultured in 21% O_2_ with or without 10 ng/mL TGFβ and 50 ng/mL HGF or cultured in 1% O_2_ were subjected to k-nearest neighbor clustering on the pcEMT gene signature into two clusters and UMAP projections were created. **(B)** Scores for gene enrichment were calculated with UCell for the epithelial and mesenchymal portions of the pcEMT gene signature, identifying one cluster enriched in epithelial genes and one enriched in mesenchymal genes. A Mann-Whitney test was performed, and the bar denotes the median. **(C)** The experimental conditions were mapped to the UMAP projection shown in (A). **(D)** Expression of the epithelial gene *CDH1* and the mesenchymal gene *VIM* were displayed on the UMAP projection from (A). **(E)** The top 10 differentially expressed genes per cluster were calculated. **(F)** Gene set variation analysis for KEGG signaling pathways was performed comparing the epithelial (E) and mesenchymal (M) clusters of the aggregated data set. **(G)** Scores for enrichment of the KEGG Hippo Signaling Pathway were calculated for the E and M clusters. A Mann-Whitney test was performed, and the bar denotes the median.

To identify regulators of EMT across the three experimental conditions, gene set variation analysis (GSVA) enrichment scores were calculated for Kyoto Encyclopedia of Genes and Genomes (KEGG) signaling pathways for a comparison of the E and M clusters. Hippo signaling was the most differentially enriched pathway (**Figure 2F**) and was preferentially active in the E cluster (**Figure 2G**). This is consistent with findings that Hippo activity promotes YAP/TAZ degradation in epithelial cells and that Hippo suppression promotes YAP/TAZ nuclear localization and expression of EMT transcription factors (26).

### Growth factors and hypoxia drive unique and shared differentially expressed genes between epithelial and mesenchymal cells

To identify common and distinct transcript features associated with EMT in response to growth factors and hypoxia, cells were separated by treatment condition prior to clustering based on the pcEMT gene signature (**Figure 3A-C**). Surveying pcEMT genes by enrichment scores for epithelial and mesenchymal genes (**Figure 3D**) or using a heatmap (**Figure 3E**), demonstrates that the clusters identified for each treatment condition are enriched in E or M genes. The proportion of cells falling into E and M clusters differed by treatment condition, as expected based on the immunofluorescence results in Figure 1A. M clusters from growth factor and hypoxic conditions were enriched in mesenchymal genes, and growth factor-treated mesenchymal cells had an overall lower expression of epithelial genes compared to hypoxic mesenchymal cells. Based on the heatmap, some genes are consistently enriched across E or M clusters regardless of treatment (e.g., *VIM* in M clusters). However, some genes are only enriched in E or M clusters for a particular condition (e.g., *AXL* in hypoxic M cells). When cells are projected in a two-PC space based on the pcEMT gene set, there is clear overlap of the three epithelial clusters but good separation for the mesenchymal clusters (**Figure 3F**), indicating that different modes of EMT lead to differential enrichment of EMT-associated genes.

**Figure 3.**
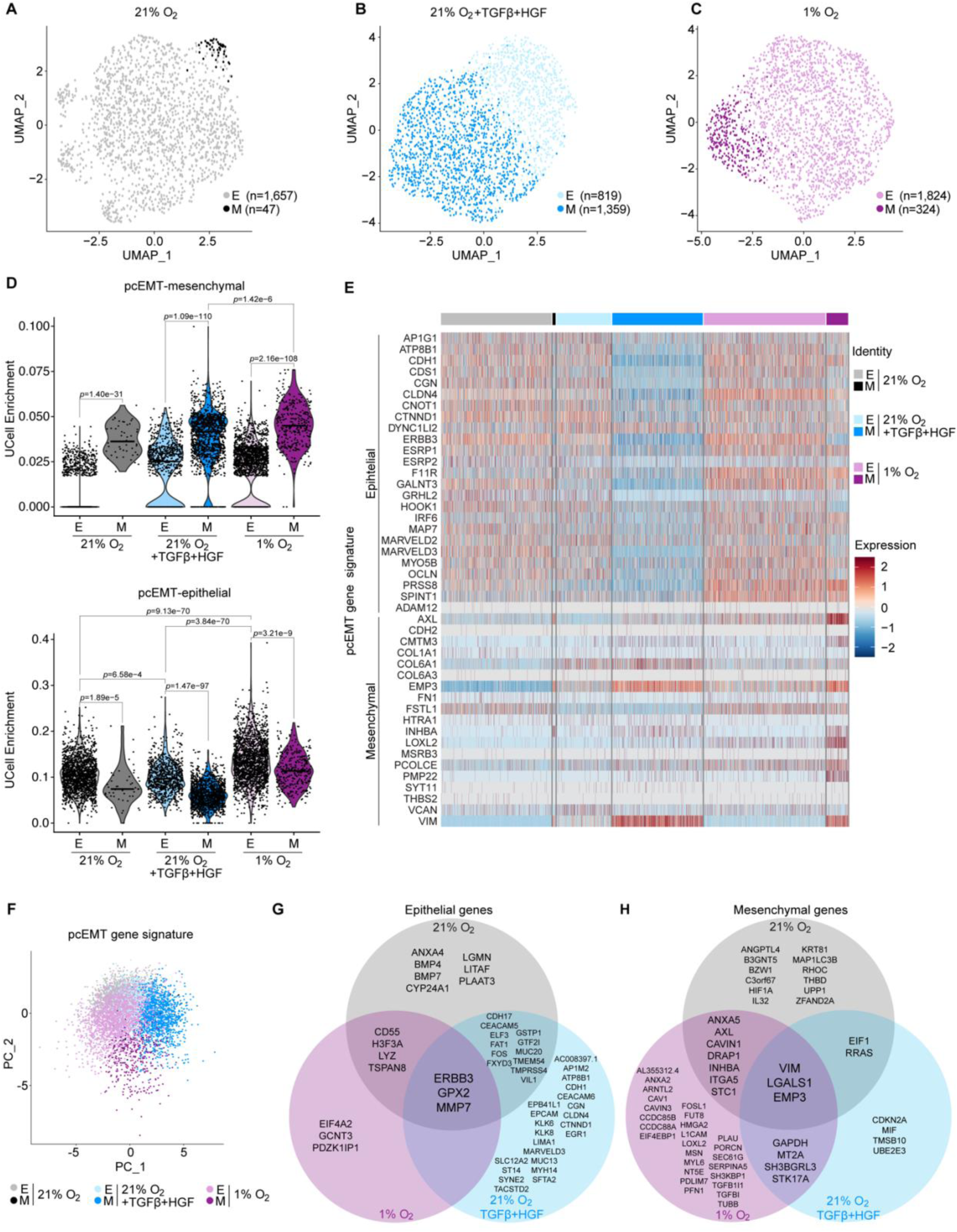
Growth factor- and hypoxia-driven EMT promote both unique and shared differential gene expression patterns. **(A-C)** HPAF-II cells cultured in **(A)** 21% O_2_, **(B)** 21% O_2_ with 10 ng/mL TGFβ and 50 ng/mL HGF, or **(C)** 1% O_2_ were subjected to k-nearest neighbor clustering on the pcEMT gene signature into two clusters and UMAP projections were created. **(D)** Scores for gene enrichment were calculated with UCell for the epithelial and mesenchymal portions of the pcEMT gene signature, identifying one clustered enriched in epithelial genes (E) and one enriched in mesenchymal genes (M). A Kruskal-Wallis test with Dunn pairwise comparisons was performed, and the bar denotes the median. **(E)** The heatmap displays the expression of the pcEMT genes for each of the E and M clusters by treatment condition. Only genes expressed in more than 50 cells are displayed. **(F)** A two-component PCA based on the pcEMT gene set features is shown for the aggregated data. Cells are color-coded by treatment condition and E/M identify based on clustering performed in panels (A)-(C). **(G,H)** The top 50 differentially expressed genes were calculated per condition between the E and M clusters, as shown in Supp Figure S3. Venn diagrams display the unique and shared **(G)** epithelial and **(H)** mesenchymal genes out of those top 50 differentially expressed per condition.

To understand the regulation of EMT in the three settings, the top 50 differentially expressed genes between the E and M clusters were identified for the three experimental conditions **(Supp Figure S3A-C)**. Of the top 50 genes per treatment, three were shared among each of the E clusters (*ERBB3*, *GPX2*, *MMP7*), and three were shared among each of the M clusters (*VIM*, *LGALS1*, *EMP3*) (**Figure 3G,H**). Further, among the top 50 differentially expressed genes, there were some genes common between just two of the E clusters (e.g., *FAT1* and *CDH17* for 21% O_2_ and growth factor-treated) or M clusters (e.g., *AXL* and *ITGA5* for 21% and 1% O_2_), as well as many that were unique to a treatment condition (e.g., *LOXL2* for 1% O_2_ M cluster). However, it is important to note that this analysis identified the top differentially expressed genes between the E and M clusters per treatment, meaning that genes appearing as unique or among just two clusters could still be altered in other conditions but not among the top 50 differentially expressed genes.

### FAT1 is enriched in epithelial cells

After surveying the results of Figure 3 and the literature, we sought to investigate and validate the functions gene products that could be hypothesized to maintain the epithelial state or promote EMT. *FAT1* enrichment was common to the 21% O_2_ and growth factor-treated epithelial clusters for the top 50 differentially expressed genes (**Figure 3G**). Direct examination of *FAT1* expression demonstrates its enrichment in epithelial cells across for all three conditions (**Figure 4A**). While *FAT1* was not among the top 50 differentially expressed genes for the hypoxic condition, *FAT1* was still significantly depleted in hypoxic M cells, with an even larger fold change than growth factor-treated E to M cells (**Figure 4A**). FAT1 suppression promotes a hybrid EMT in squamous cell carcinoma (27), and FAT1 activates the Hippo pathway in head and neck squamous cell carcinoma (18). In the aggregated data set, *FAT1* expression positively correlates with Hippo pathway enrichment (**Figure 4B**). Based on this, we probed the relevance of *FAT1* and its gene product in determining the epithelial phenotype.

**Figure 4.**
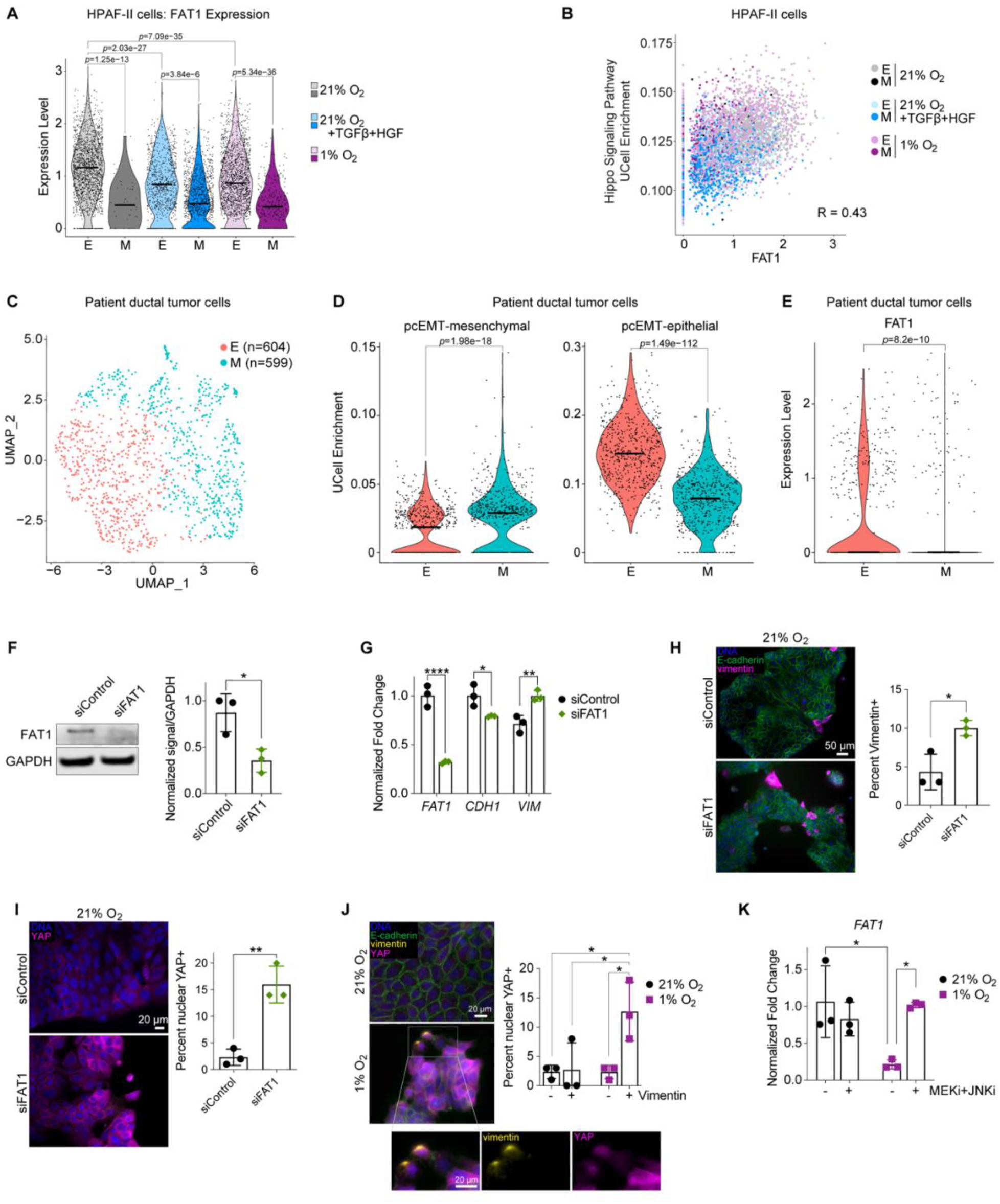
FAT1 expression is higher in epithelial cells. **(A)** The expression of *FAT1* per cell is displayed per E/M cluster per condition. A Kruskal-Wallis test with Dunn pairwise comparisons was performed, and the bar denotes the median. **(B)** The scatter plot displays the UCell enrichments for the KEGG Hippo Signaling Pathway and *FAT1* expression per cell for the aggregated data as annotated by E/M clusters per condition. The Pearson correlation was calculated. **(C)** scRNA-seq from patient tumors (28) was subjected to the same analysis as with HPAF-II cells. A UMAP projection displays k-nearest neighbor clustering on the pcEMT gene signature into two clusters. **(D)** Scores for gene enrichment were calculated with UCell for the epithelial and mesenchymal portions of the pcEMT gene signature, identifying one cluster enriched in epithelial genes (E) and one enriched in mesenchymal genes (M). A Mann-Whitney test was performed, and the bar denotes the median. **(E)** The expression of *FAT1* per cell is displayed comparing the E and M cluster of the patient tumor data. A Wilcoxon test was performed, and the bar denotes the median. **(F)** HPAF-II cells were transfected with siRNA targeting FAT1 or control siRNA and cultured at 21% O_2_ for 120 hr. Lysates were analyzed by immunoblotting for the indicated proteins. *n* = 3, t test. **(G)** qRT-PCR was performed for the indicated transcripts on RNA isolated from HPAF-II cells treated as described in (F). *CASC3* was used as a control gene for normalization. *n* = 3, two-way ANOVA with Sidak’s multiple comparisons test comparing FAT1 to control siRNA for each transcript. **(H,I)** HPAF-II cells were treated as in (F) and immunofluorescence microscopy was performed for the indicated proteins. *n* = 3, t test. **(J)** HPAF-II cells were cultured in 21% or 1% O_2_ for 120 hr and immunofluorescence microscopy was performed for the indicated proteins. *n* = 3, two-way ANOVA with Tukey’s multiple comparison test. **(K)** HPAF-II cells were cultured in 21% or 1% O_2_ with 1 μM CI-1040 (MEKi) and 10 μM SP600125 (JNKi), or DMSO for 120 hr, with inhibitors replenished every 48 hr. RNA was extracted, and qRT-PCR was performed for *FAT1*, with *CASC3* as a control gene for normalization. *n* = 3, two-way ANOVA with Tukey’s multiple comparison test. * *p* < 0.05, ** *p* < 0.01, *** *p* < 0.001, **** *p* < 0.0001

To analyze the role *FAT1* in the intact tumor setting, we used a published scRNA-seq data set of six PDAC patient tumors (28). Limiting the analysis to ductal cells only, the pcEMT signature was used to create two clusters (**Figure 4C**), which were annotated as E and M based on their differential enrichment for mesenchymal and epithelial genes **(Figure 4D, Supp Figure S4)**. Ductal cells in the epithelial cluster displayed greater *FAT1* expression than those in the mesenchymal cluster (**Figure 4E**), consistent with the findings in HPAF-II cells.

To probe these inferences functionally *in vitro*, we knocked down *FAT1* in HPAF-II cells (**Figure 4F**). *FAT1* depletion suppressed *CDH1* and promoted *VIM* transcript expression (**Figure 4G**) and promoted vimentin protein expression (**Figure 4H**). FAT1-depleted cells also displayed increase nuclear accumulation of YAP (**Figure 4I**), consistent with prior reports that FAT1 antagonizes YAP signaling by assembling the Hippo pathway (18). YAP nuclear localization was most common in vimentin-positive hypoxic cells (**Figure 4J**). Thus, FAT1 depletion may promote YAP nuclear localization and resultant EMT.

We further tested whether the relationship between the current findings and our prior work demonstrating ERK and JNK signaling as required for hypoxia-mediated EMT (3). Combined MEK and JNK inhibition rescued *FAT1* expression in hypoxic culture (**Figure 4K**), confirming key roles for these kinases in regulating *FAT1* as a feature of the epithelial cell state.

### AXL regulates hypoxia-mediated EMT

The striking result in Figure 3E suggesting that *AXL* is primarily enriched in the hypoxic M cluster, and not in growth factor-treated M cells motivated investigation, motivated testing of AXL’s function in hypoxia. In breast cancer, AXL can be a driver of EMT potentially through NFκB signaling (29), but AXL can also be a result of a mesenchymal transition with vimentin being required for AXL expression (21) . Therefore, it is unclear if AXL plays a supporting role in EMT or if it is a consequence of EMT. While little is known about the relationship between AXL and EMT in PDAC, AXL inhibition promotes PDAC chemosensitivity (30), making AXL an enticing target to explore.

To expand on the observation in Figure 3E, we confirmed that *AXL* is significantly enriched in hypoxic mesenchymal cells (**Figure 5A**). To determine if this also occurs in the intact tumor microenvironment, we clustered ductal PDAC tumor cells from a published patient tumor scRNA-seq data set using the Hallmark Hypoxia gene set (**Figure 5B**), which identified a cluster enriched in Hallmark Hypoxia and HIF-target genes, denoted as “HYP+” (**Figure 5C**). HYP+ cells were then clustered using the pcEMT gene set features (**Figure 5D**), which produced clusters enriched in E or M genes (**Figure 5E**). Within the pcEMT gene set, *AXL* was the most consistently differentially regulated gene between the E and M clusters (**Figure 5F**). Consistent with the analysis of HPAF-II cells, *AXL* was preferentially enriched in hypoxic M cells (**Figure 5G**). Interestingly, AXL inhibition prevented EMT in response to hypoxia but not in response to growth factors, suggesting a context-specific role for AXL in driving EMT (**Figure 5H**).

**Figure 5.**
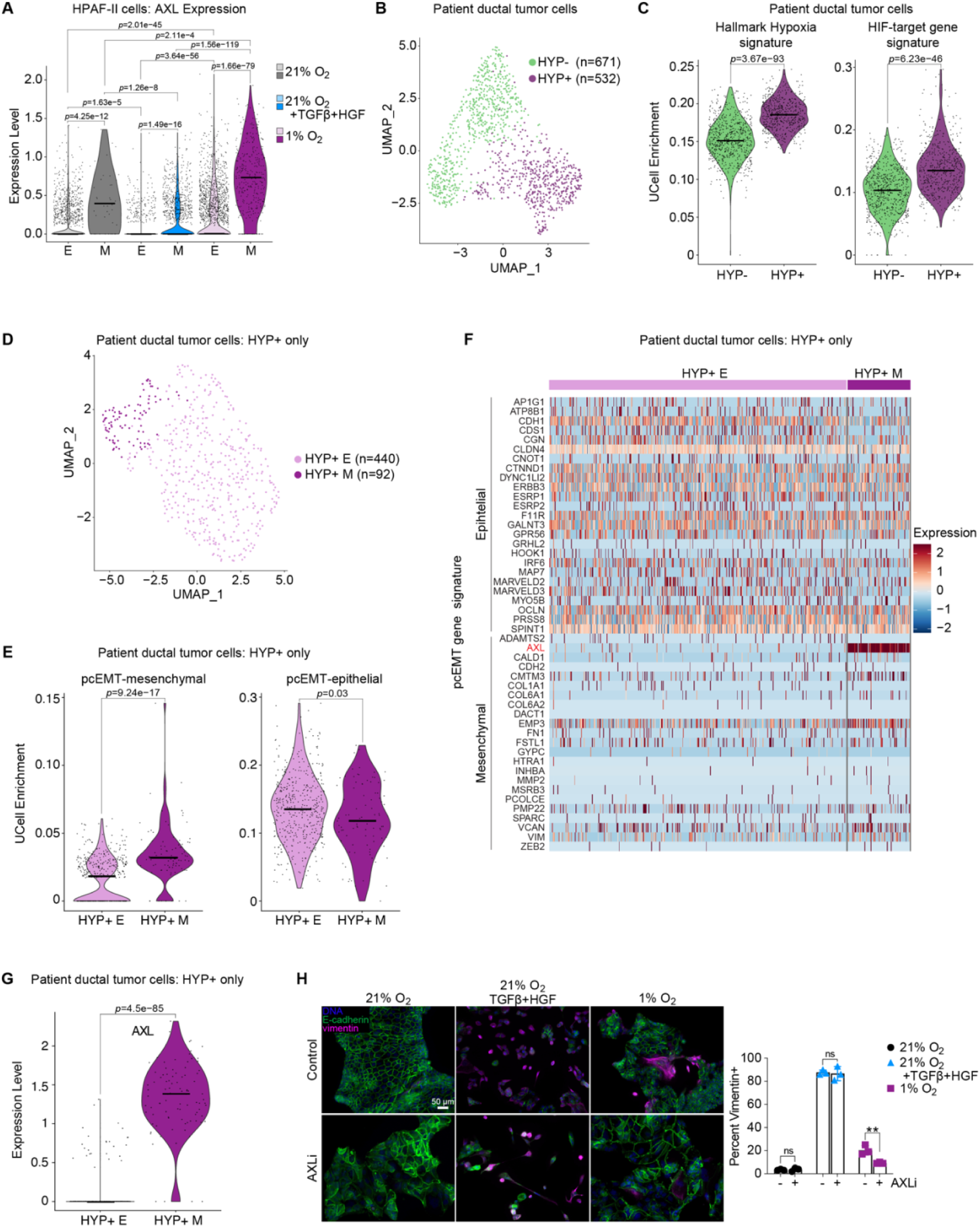
AXL is preferentially enriched in hypoxic mesenchymal cells. **(A)** The expression of *AXL* per cell is displayed per E/M cluster per condition. A Kruskal-Wallis test with Dunn pairwise comparisons was performed, and the bar denotes the median. **(B)** A UMAP projection displays k-nearest neighbor (kNN) clustering on scRNA-seq from patient tumors (28) on the Hallmark Hypoxia gene set into two clusters. **(C)** Scores for gene enrichment were calculated with UCell for the Hallmark Hypoxia gene signature and HIF-target gene signature, identifying one clustered enriched in hypoxic genes (HYP+) compared to the other cluster (HYP-). A Mann-Whitney test was performed, and the bar denotes the median. **(D)** The HYP+ cluster from (B) was then subjected to kNN clustering using the pcEMT gene signature, as previously done with the HPAF-II data set. **(E)** Scores for gene enrichment were calculated with UCell for the epithelial and mesenchymal portions of the pcEMT gene signature, identifying one cluster enriched in epithelial genes (E) and one enriched in mesenchymal genes (M). A Mann-Whitney test was performed, and the bar denotes the median. **(F)** The heatmap displays the expression of the pcEMT genes for each of the E and M clusters for the HYP+ patient tumor ductal cells. **(G)** The expression of *AXL* per cell is displayed comparing the E and M cluster of the HYP+ patient tumor data. A Wilcoxon test was performed, and the bar denotes the median. **(H)** HPAF-II cells were cultured in 21% O_2_ with or without 10 ng/mL TGFβ and 50 ng/mL HGF or cultured in 1% O_2_, with 40 nM dubermatinib (AXLi) and or DMSO for 120 hr. Immunofluorescence microscopy was performed for the indicated proteins. *n* = 3, two-way ANOVA with Sidak’s multiple comparison test. ** *p* < 0.01

Given the functional relevance of AXL, we sought to understand the protein expression of AXL in response to the EMT drivers. AXL intensity was greatest in hypoxic cells, but only nuclear AXL was increased in vimentin-positive cells. Increased nuclear AXL was observed in vimentin-positive cells from TGFβ+HGF or hypoxic treatment, with hypoxic mesenchymal cells displaying the highest overall nuclear AXL intensity (**Figure 6A**). In some settings, nuclear AXL represents a proteolytic cleavage product of the full-length receptor containing a nuclear localization sequence (31), but there is still limited understanding for the role of nuclear AXL. Lysates of HPAF-II cells do not suggest the presence of a lower molecular weight cleavage in hypoxic culture though (**Figure 6B**), despite the higher nuclear intensity by immunofluorescence microscopy. Given that YAP promotes *AXL* transcription (20), we tested for a correlation between YAP and AXL and found that nuclear AXL intensity was greatest in hypoxic cells displaying nuclear YAP (**Figure 6C**). These results suggest a role of nuclear AXL in promoting EMT in mesenchymal hypoxic cells. Based on the aggregated analyses presented, we propose a mechanism wherein FAT1 loss inhibits the Hippo pathway and allows for YAP nuclear localization and resulting AXL expression, which promotes EMT.

**Figure 6.**
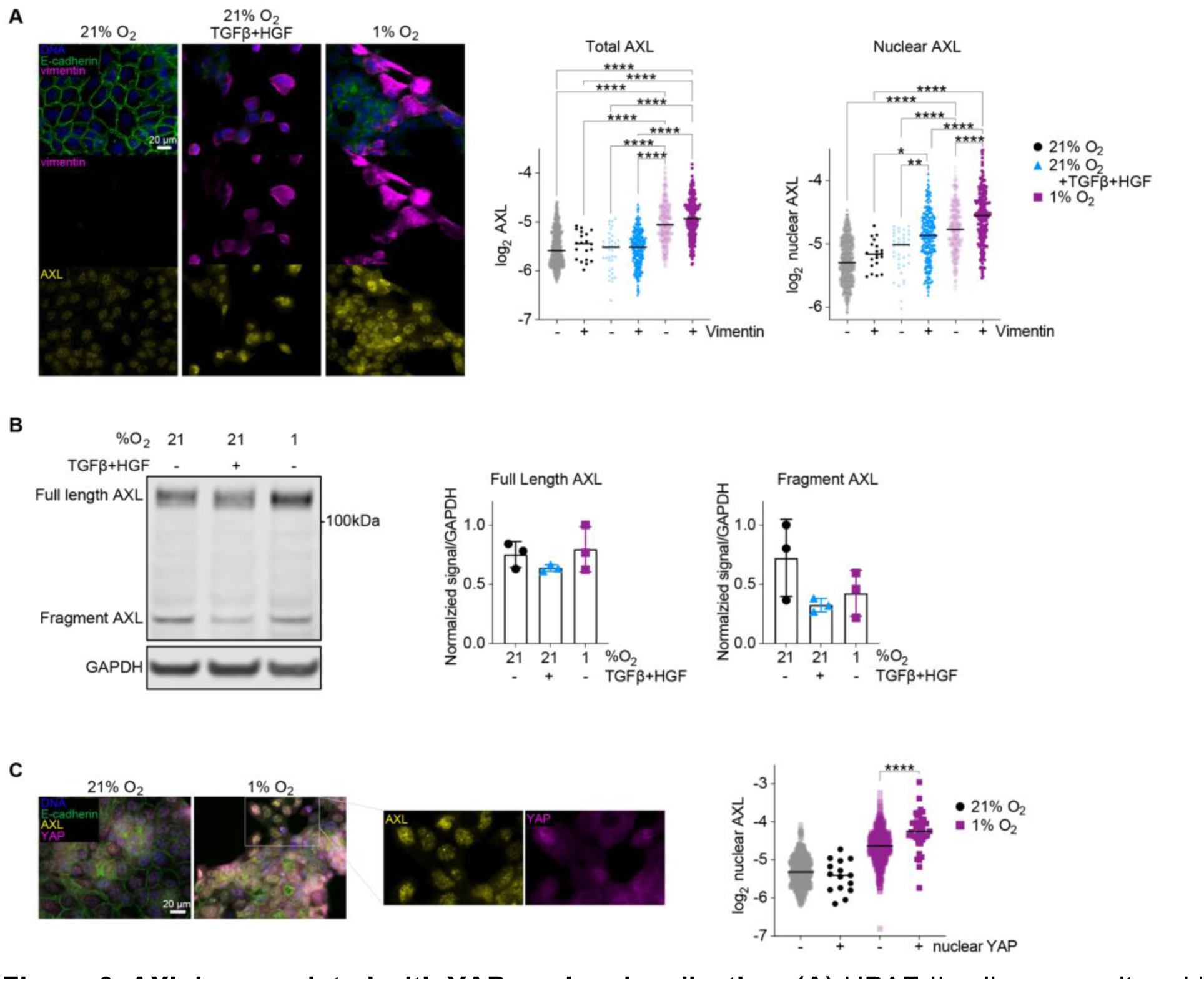
AXL is associated with YAP nuclear localization. **(A)** HPAF-II cells were cultured in 21% O_2_ with or without 10 ng/mL TGFβ and 50 ng/mL HGF or cultured in 1% O_2_ for 120 hr. Immunofluorescence microscopy was performed for the indicated proteins. *n* = 3, one-way ANOVA with Tukey’s multiple comparisons test. **(B)** HPAF-II cells were cultured as in (A). Cells lysates were analyzed by immunoblotting for the indicated proteins. **(C)** HPAF-II cells were cultured in 21% or 1% O_2_ for 120 hr and immunofluorescence microscopy was performed for the indicated proteins. *n* = 3, one-way ANOVA with Tukey’s multiple comparisons test. * *p* < 0.05, ** *p* < 0.01, *** *p* < 0.001, **** *p* < 0.0001

## DISCUSSION

In this study, we utilized scRNA-seq to understand heterogenous EMT transcriptional regulation in populations of pancreas cancer cells in response to different drivers of the mesenchymal transition. We identified both common and unique modes of EMT regulation at play between growth factor and hypoxia conditions. FAT1 was enriched across epithelial cells regardless of the treatment condition, whereas AXL played a context-dependent role in hypoxia-mediated EMT only. Previous work found that *FAT1* deletion promoted a hybrid EMT through YAP nuclear localization and ZEB1 expression in squamous cell carcinoma (27), and that FAT1 prevents EMT via suppression of the MAPK/ERK pathway in esophageal squamous cell cancer (32). Additionally, to account for the hypoxia-specific enrichment of AXL, studies have found that HIF-1⍺ and HIF-2⍺ directly promote AXL expression, which promotes SRC activity (33). We have previously established a role for SRC-dependent MAPK/ERK-signaling in regulating hypoxia-mediated EMT (3), which unifies the present study and previous findings on FAT1 and AXL to provide a mechanism for hypoxia-driven EMT in PDAC.

Aside from its role in EMT, AXL is involved in cell survival, angiogenesis, and immune response in pancreas cancer (34). AXL-deficient mice bearing pancreas tumors have increased survival, more differentiated tumor histology, and decreased metastasis, making AXL an attractive therapeutic target (35). Our results suggest a potentially interesting role for nuclear AXL in correlation with EMT. Nuclear AXL has been observed in various carcinomas, including schwannoma, melanoma, and mesothelioma (36–39); however, there is still more to interrogate regarding the function of nuclear AXL, especially in relation to EMT. One study identified in mesothelioma that there was nuclear colocalization of AXL and p53, and that AXL can bind to the *TP53* promoter to suppress expression (39). Several AXL inhibitors are in clinical trials for advanced solid tumors, colorectal cancer, non-small cell lung cancer, and other diseases (ClinicalTrials.gov). However, only one clinical trial has tested AXL inhibitors in pancreas cancer, in combination with chemotherapy, and that study was terminated (NCT03649321; ClinicalTrials.gov). There has yet to be another study initiated for AXL inhibition in PDAC. Given the bias for AXL expression in hypoxic mesenchymal cells, more success could be achieved from AXL inhibition by targeting AXL-driven EMT in cooperation with other therapies. Supporting the notion of an important role for AXL in PDAC pathogenesis, soluble AXL is a useful biomarker for classifying patients as healthy, tumor-burdened, or having chronic pancreatitis (40). Further, a current clinical trial in PDAC detects AXL-positive circulating tumor cells (CTCs) through real-time liquid biopsy to eventually correlate survival with the presence of AXL-positive CTCs (NCT05346536; ClinicalTrials.gov). Therefore, although AXL inhibition is still being investigated, detection of AXL could provide critical diagnostic insight in PDAC.

While our study focuses on EMT transcriptional regulation, post-translational and epigenetic modifications may also play important roles in determining EMT heterogeneity. The activity of different EMT transcription factors can be dependent on the context and stage in transition (41). Further, the localization and turnover of EMT transcription factors, including Snail, Slug, Twist, and ZEB1, are regulated by acetylation, phosphorylation, and ubiquitination (42,43). Mutations and post-translational modifications can also influence the activity of EMT-regulating signaling pathways, such as mutant KRAS promoting EMT in a MEK-dependent manner (44) and EGF-stimulated SHP2 activity augmenting EMT (7). Further, the Hippo pathway constrains YAP through phosphorylation-dependent degradation via LATS1/2 (45,46). In specific settings, histone methylation and acetylation marks are required for EMT, including histone 3 lysine 4 acetylation (H3K4Ac), histone 3 lysine 4 di-methylation (H3K4me2), histone 3 lysine 27 tri-methylation (H3K27me3), and histone 3 lysine 36 di-methylation (H3K36me2) (47–49). At least one of these marks (i.e., H3K36me2) is induced by both EMT-driving growth factors and hypoxia (47–49). Here, our analysis nominates H3 Histone Family 3A (*H3F3A*) as enriched in cells cultured in 21 or 1% O_2_, potentially pointing to a context-specific role for H3F3A. Mass cytometry and high-content, multi-channel immunofluorescence imaging will be important tools moving forward for discovering the potential supporting role of post-translational modifications in explaining EMT heterogeneity.

Given that mesenchymal cells are more resistant to chemotherapy (50), heterogeneity of EMT poses a significant challenge to treatment. One of the ways to combat this across carcinomas could be molecular subtyping to identify patients that will likely be more responsive (51). In PDAC, there are ongoing studies to classify patient tumors as classical or basal-like, since classical tumors are more responsive to the chemotherapy FOLFIRINOX (52). However, as identified through single-cell measurements in this study and others, there is significant heterogeneity in EMT between cells within a cell line and within a tumor. Since mesenchymal cells are chemoresistant (53), the existence of heterogenous EMT states creates a subpopulation of cells that are less chemoresponsive and likely to remain after treatment (54). A means to combat this heterogeneity could be to use small molecule drugs as neoadjuvants to antagonize EMT prior to conventional chemotherapy (55). Due to the heterogeneity of EMT and the regulatory processes that control it, identifying the most appropriate molecularly targeted drugs must leverage measurement and analysis techniques that accommodate cell-to-cell variability.

## METHODS

### Cell culture

HPAF-II cells (Carl June, University of Pennsylvania) were cultured in RPMI with 10% fetal bovine serum (FBS), 1 mM L-glutamine, 100 units/mL penicillin, and 100 μg/mL streptomycin. The Genetic Resources Core Facility at the John Hopkins University School of Medicine authenticated the HPAF-II cells by performing short tandem repeat profiling via GenePrint 10 (Promega) and comparing to the ATCC database. HPAF-II cells were assessed for mycoplasma using the MycoAlert PLUS Detection Kit (Lonza). Cells were cultured in a Thermo Scientific Forma Steri-Cycle i160 incubator at 5% CO_2_ and 37°C for normal culture. For hypoxic culture at 1% O_2_, cells were grown in a Tri-Gas version of the same incubator, which allows for N_2_ to displace O_2_. The lower limit of the incubator is 1%. Following plating, cells cultured under normal 21% O_2_ conditions for 16 hr then either maintained in 21% or transferred to the tri-gas incubator.

### Growth factors and inhibitors

Recombinant human TGFβ and HGF (Peprotech) were used at 10 ng/mL and 50 ng/mL, respectively. During treatment, complete medium with growth factors was replenished every 48 hr. The AXL inhibitor dubermatinib (MedChemExpress) was used at 40 nM, MEK inhibitor CI-1040 (LC Laboratories) was used at 1 μM, and JNK inhibitor SP600125 (LC Laboratories) was used at 10 μM. Stocks of all inhibitors were prepared in DMSO.

### Antibodies

Antibodies against vimentin (Santa Cruz Biotechnology, sc-373717), E-cadherin (clone ECCD2, Invitrogen, 13-1900), FAT1 (BiCell, 50501), AXL (Cell Signaling Technology, #8661), YAP (Santa Cruz Biotechnology, sc-101199), and GAPDH (Santa Cruz Biotechnology, sc-32233) were used. For a nuclear stain, Hoechst 33342 (Invitrogen, H1399) was used.

### Coverslip immunofluorescence

Cells were cultured on 18-mm glass coverslips in 6-well plates. At the conclusion of an experiment, cells were fixed with 4% paraformaldehyde in PBS for 20 min and then permeabilized with 0.25% Triton-X 100 in PBS for 5 min. Primary antibodies were added to coverslips in a humidified chamber overnight at 4°C. Coverslips were washed five times with 0.1% Tween 20 in PBS, then incubated for 1 hr at 37°C in a humidified chamber with Alexa Fluor secondary antibodies and Hoechst nuclear stain. All antibodies were diluted in Intercept Blocking Buffer (Licor, 927-60001). Following the staining and washing steps, coverslips were mounted on glass slides with ProLong Gold Antifade Mountant.

### Fluorescence microscopy and automated image analysis

Coverslips were imaged on a Zeiss Axiovert Observer.Z1 fluorescence microscope, using a 20 or 63× objective and ZEN image processing software to produce .czi files. Within a particular experiment, across replicates and conditions, imaging was performed using the same exposure times and image settings. Four frames were imaged at random for each biological replicate, yielding at least 1000 cells per replicate. For image analysis, CellProfiler v3.1.9 (Broad Institute) was used to quantify signal intensity and localization (56–58). Discrete cells were identified based on the nuclear stain as the primary object for the analysis pipeline. For percent-positive measurements, a threshold was set based on a negative control consisting of a sample stained with a secondary antibody only. All cells were subject to the same threshold, and a percentage was calculated based on the number of cells with signal above background relative to the total cell number. For intensity measurements, the mean intensity per object was measured. For nuclear measurements, the nuclear stain was used to restrict the signal to the nuclear domain.

### Western blotting

Cells were lysed using a standard cell extraction buffer (Invitrogen, FNN0011) supplemented with protease and phosphatase inhibitors (Sigma-Aldrich, P8340, P5726, P0044). Crude lysates were centrifuged at 14,000 rpm for 10 min at 4°C, and supernatants were retained as clarified lysates. Total protein concentration was measured with a micro-bicinchoninic acid (BCA) assay (Pierce). Equal protein amounts were mixed with 10× NuPAGE reducing agent, 4X LDS sample buffer and MilliQ water. Samples were heated for 10 min at 100°C and loaded onto a 1.5 mm NuPAGE gradient (4-12%) gel (Invitrogen, NP0336BOX). After electrophoresis, gels were transferred to a 0.2 μm nitrocellulose membrane using the TransBlot Turbo Transfer System (BioRad). Membranes were blocked with Intercept Blocking Buffer (IBB; Licor, 927-60001) for 1 hr on an orbital shaker. Primary antibodies were diluted at 1:1000 in IBB, then incubated with membranes overnight at 4°C. Membranes were washed with 0.1% Tween-20 in PBS by shaking three times for 5 min each. Secondary antibodies were diluted 1:10,000 in IBB and incubated with shaking for 2 hr at room temperature. Membranes were washed as previously described with 0.1% Tween-20 in PBS. Membranes were imaged on a LiCor Odyssey. Image Studio software was used for band densitometry.

### Single-cell RNA-sequencing

scRNA-seq was performed by the UVA Genome Analysis and Technology Core. HPAF-II cells were cultured in 6-well plates with three biological replicates per condition, with approximately one million cells per replicate at the conclusion of the experiment. For sequencing, three biological replicates were pooled together prior to preparing the samples. Cells were prepared using the 10× Genomics® Sample Preparation Demonstrated Protocol, *Single Cell suspensions from Cultured Cell Lines for Single Cell RNA Sequencing* (Manual CG00054 Rev B). After confirming cell viability and single cells, Next Generation Sequencing (NGS) Library preparation was performed for 3’ RNA gene expression. For quality control of the library, a Qubit Fluorometer (Thermo Fisher), TapeStation D5000 HS system (Agilent), and MiSeq – 300 cycle nano (Illumina) were used. Sequencing was performed using Illumina NextSeq^TM^2000 with P2 – 100 cycle v3 to allow for 25,000 reads per cell.

### Gene sets and signatures

The pan-cancer EMT signature (25) was used as the primary feature set for clustering based on EMT markers. The HIF target signature was used as previously published (24). Hallmark gene sets were obtained from the Molecular Signatures Database, including Hallmark Hypoxia and Hallmark EMT (59). Gene sets from the Kyoto Encyclopedia of Genes and Genomes (KEGG) (60,61) were accessed within R using the *clusterProfiler* package.

### Pre-processing of scRNAseq data

10× genomics data was aligned to the human genome and filtered to contain only cells with barcodes using CellRanger by the UVA Bioinformatics Core. This created a filtered dataset of 1910 (normoxic), 2493 (TGFβ+HGF), and 2441 (hypoxic) cells. Software and R packages used are cited in **Supp Table S1**. The code used to analyze the data can be accessed here: https://github.com/lazzaralab/Brown-et-al_PDAC-scRNAseq. The *Seurat* R package was used to filter, normalize, scale, and cluster the data. The data was filtered to remove low-quality cells and doublets, by filtering out cells that express too few or too many genes and excessive mitochondrial gene counts. The R package *scRNABatchQC* (62) was used to determine the threshold for these metrics. Cells that had greater than 1882 genes, fewer than 8500 genes, and less than 15 percent mitochondrial genes were retained for analysis. After quality control steps, a dataset of 1703 (normoxic), 2178 (TGFβ+HGF), and 2148 (hypoxic) cells was available for analysis. Using *Seurat*, the data was normalized by log-transforming and scaled for the mean expression to be 0 and the variance across cells to be 1.

### Principal components analysis and clustering

To reduce the dimensionality of the data, we performed principal component analysis (PCA) for either all features or a subset of features (i.e., pcEMT gene set, HIF-target gene set, or Hallmark Hypoxia gene set, as noted in the results). For analysis including all genes, the number of principal components (PCs) to include was determined by calculating 1) the point where the PCs only contribute 5% of standard deviation and the PCs cumulatively contribute 90% of the standard deviation, and 2) the percent change in variation between consecutive PCs is less than 0.1%, and using the lower value of 1) and 2) for the number of PCs to include in downstream analysis (as adapted from the Harvard Chan Bioinformatics Core). The *Seurat* package was used to employ k-nearest neighbor (kNN) clustering on a PCA projection of the data using the optimal dimensionality determined as described above. For analysis including all genes, the optimal resolution for the number of clusters was determined by calculating the silhouette score (as adapted from Roman Hillje), and the resolution was constrained when restricting the data to two clusters for comparison of E and M. Using *Seurat*, a UMAP projection was employed to visualize the clusters created from kNN clustering.

### Gene expression and gene set enrichment

The *Seurat* package was used to calculate and visualize differential gene expression between the clusters, to visualize gene expression by violin plots, and for scatter plots. The R package *Escape* was used for gene set enrichment analysis (GSEA) calculations for individual cells. Per cell enrichment values were calculated using ssGSEA, as previously described (63). Significant differences between clusters were determined by performing ANOVA. Enrichment plots for the Response to Hypoxia were performed by calculating a mean rank order for a gene set using ssGSEA across each condition. The R package *UCell* was used for scoring gene signatures based on the Mann-Whitney U statistic (64). Gene set variation analysis (GSVA) was utilized to determine the enrichment of Kyoto Encyclopedia of Genes and Genomes (KEGG) signaling pathways using the R packages *GSVA* and *limma* for statistical testing. GSVA scores were calculated and fit to a linear model, followed by empirical Bayes statistics for differential enrichment analysis, as previously described [as adapted from Roman Hillje (65)]. The log-odds that the gene set is differentially enriched was displayed.

### siRNA-mediated knockdowns

*Silencer^TM^* Select siRNA against FAT1 s5033 (Thermo Fisher, 4392420) and *Silencer^TM^*Select negative control siRNA (Thermo Fisher, 4390843) were used with Lipofectamine RNAiMAX (Thermo Fisher) per manufacturer recommendations.

### Quantitative reverse transcription PCR (qRT-PCR)

RNA was extracted using the RNeasy Kit (Qiagen, #74104) and reverse transcribed using High-Capacity cDNA Reverse Transcription Kit (Applied Biosciences, #4368814). qRT-PCR was performed using PowerUp SYBR Green (Applied Biosciences, #A25741) per manufacturer protocol using a QuantStudio3 system (Applied Biosystems). Measurements were analyzed with the ddCt method (66). Data are displayed as a normalized fold changes, using *CASC3* as a housekeeping gene. Primer sequences are provided in **Supp Table S2**.

### Statistical analyses for experimental studies

Prism 9 for macOS was used for all experimental statistical analyses. Figure captions include details on the analyses conducted. For two-way ANOVA, the post hoc test was chosen based on the comparison made, with Tukey’s multiple comparisons used when considering all conditions and Sidak’s multiple comparisons used when considering specific comparisons within a larger dataset.

## ACKNOWLEDGEMENTS

We thank Dr. David Tuveson (Cold Springs Harbor Laboratory) for sharing annotated tumor scRNAseq data. We acknowledge Dr. Katia Sol-Church at UVA Genome Analysis and Technology Core for assisting in experimental design and performing the scRNA-sequencing on HPAF-II cells. We thank Drs. Ben Stanger (University of Pennsylvania) and Jason Pitarresi (University of Massachusetts) for helpful technical discussions.

## FUNDING STATEMENT

This work was supported by NCI U01 CA243007 (MJL), an NSF Graduate Research Fellowship (BAB), the NIH Cancer Training Program 5T32CA009109 at UVA, and UVA Cancer Center Support Grant NCI P30CA044579.

## CONFLICTS OF INTEREST

The authors declare no conflicts of interest.

## DATA ACCESS STATEMENT

The authors agree to make scRNA-sequencing data publicly available at the time of publication. Code for the analysis is available as described in *Methods*.

## Notes

### Competing Interest Statement

The authors have declared no competing interest.

