## Supplemental Material for "Single-cell RNA sequencing reveals microenvironment context-specific routes for epithelial-mesenchymal transition in pancreas cancer cells"

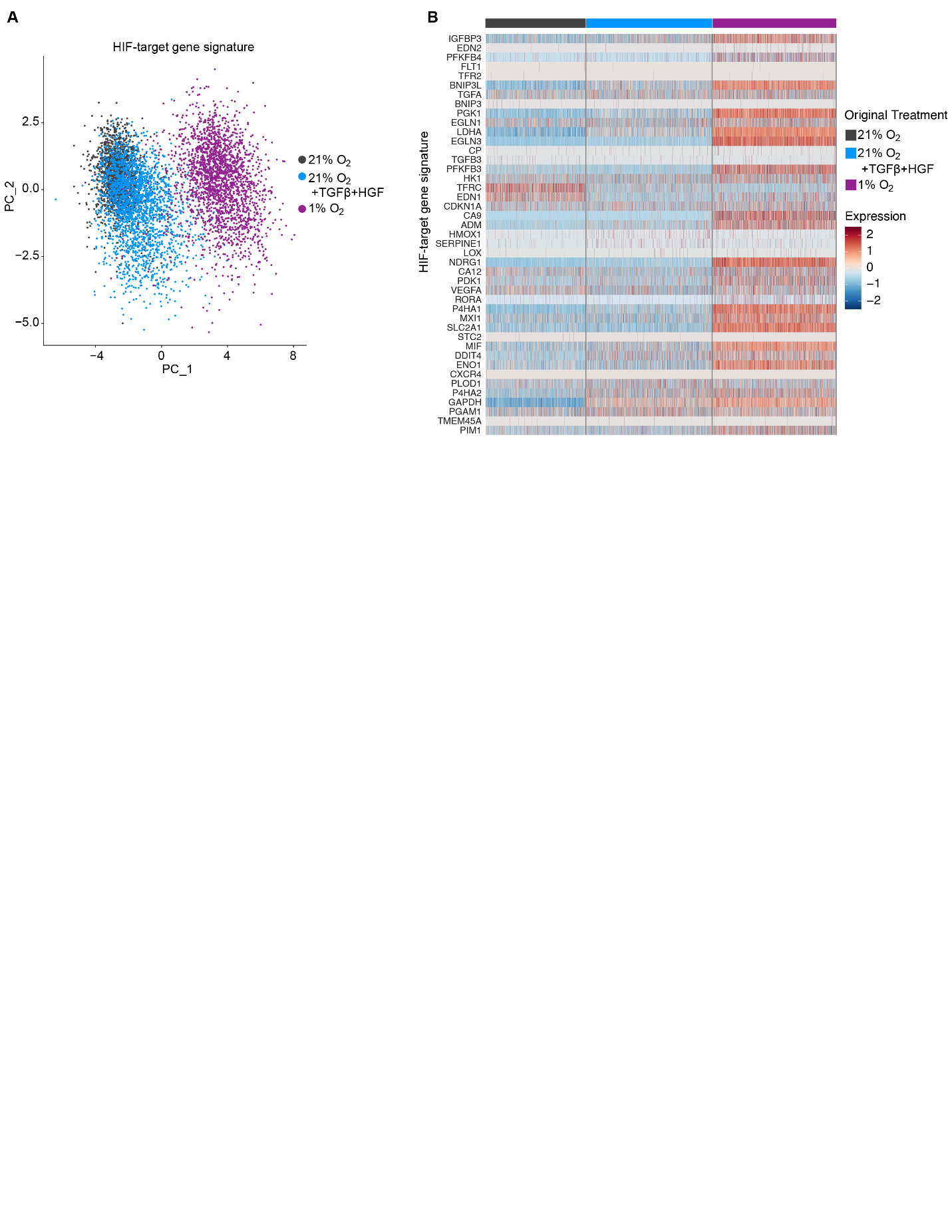

**Supp Figure S1. Hypoxic cell culture promoted expression of hypoxia-associated genes. (A)** PCA on the HIF-target gene set was computed for the aggregated data as mapped to the experimental conditions. **(B)** The heatmap displays the expression of the HIF-target genes with annotation of the experimental conditions.

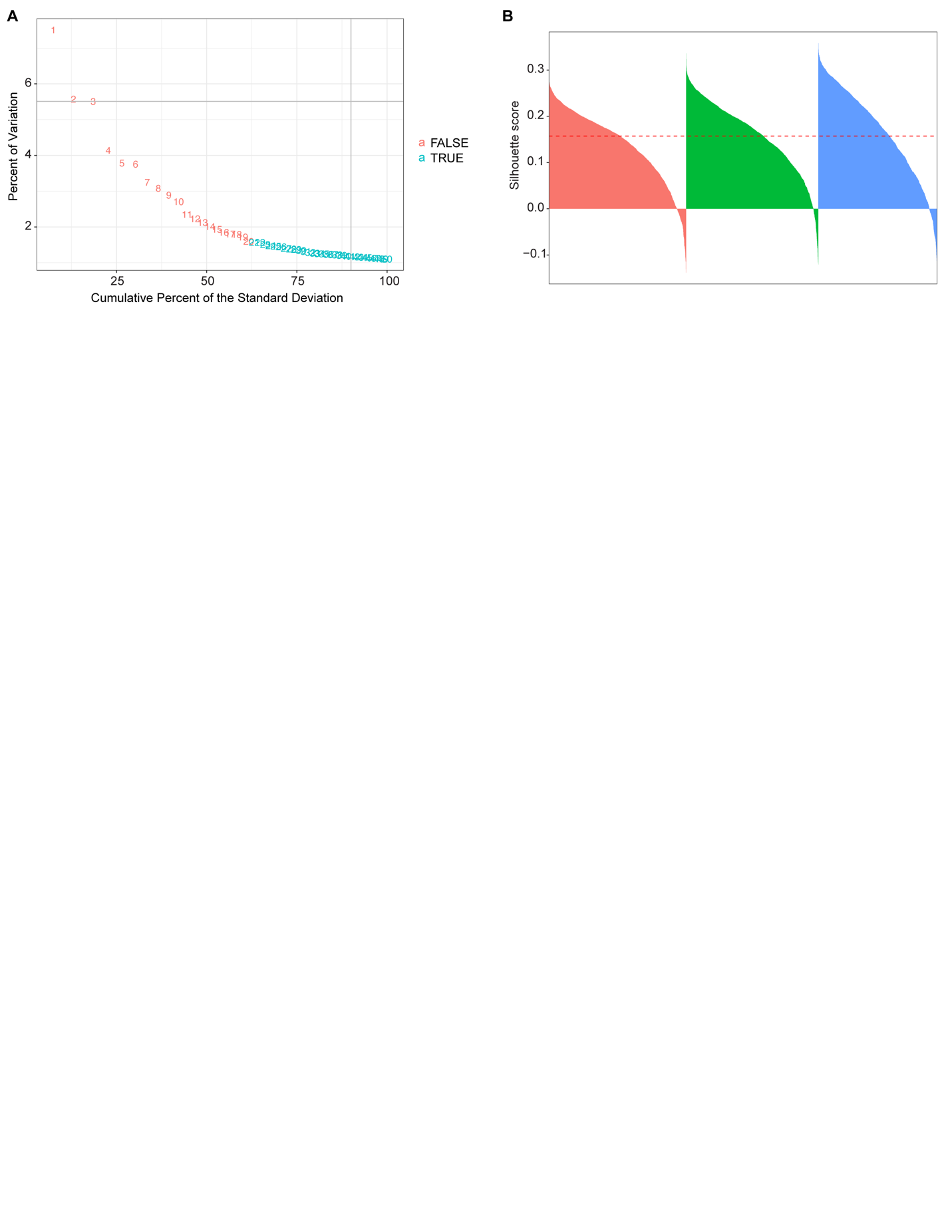

**Supp Figure S2. Optimal parameters were chosen for clustering on all genes. (A)** An elbow plot showing the percent variation and cumulative percent for each principal component used for clustering in Figure 1B. Plotted numbers refer to the principal component (PC) number. Red and blue numbers indicate the change in percent variation to the next PC is greater than or less than 0.1%, respectively. **(B)** Silhouette scores are shown per cell by cluster for the optimal number of k-nearest neighbor clusters based on the aggregated data set.

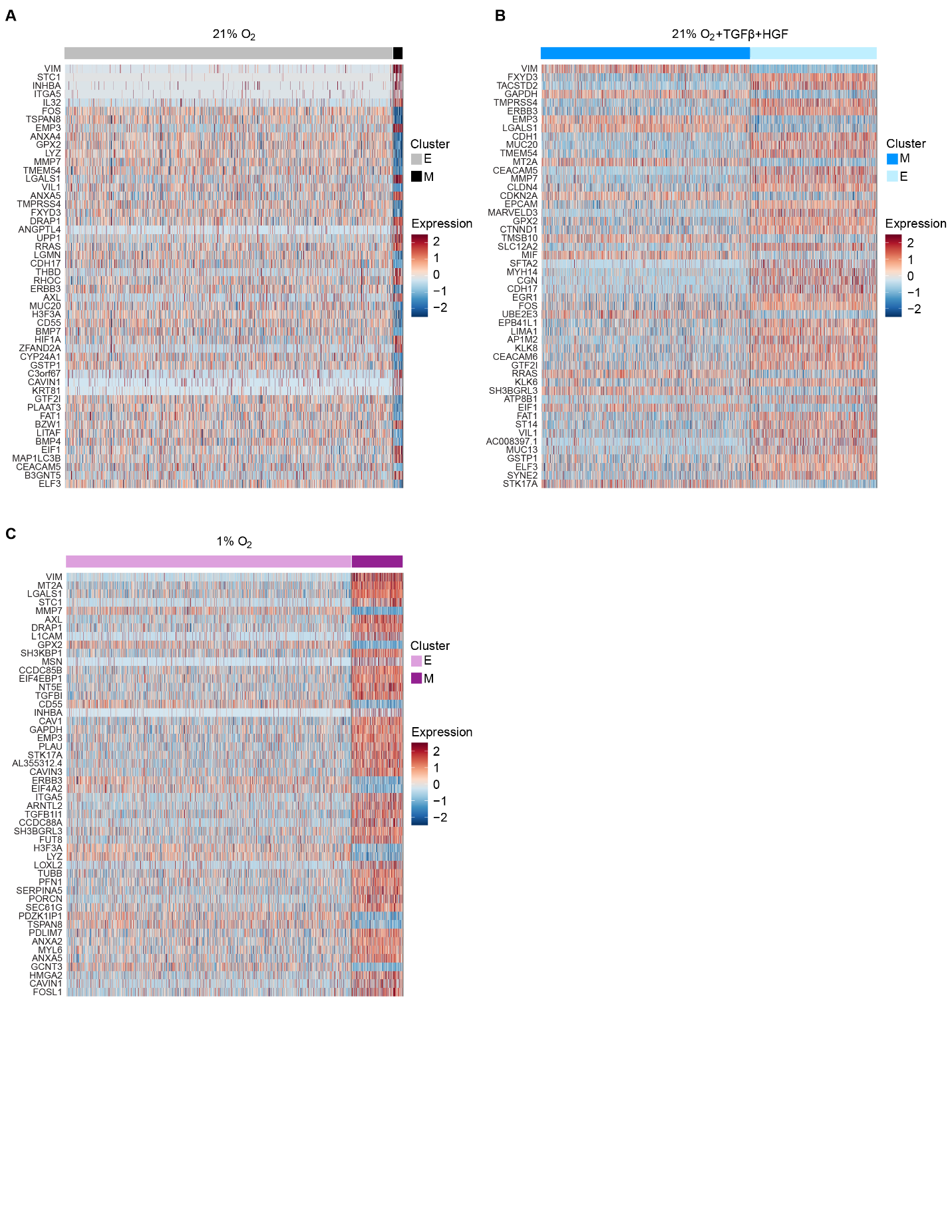
**Supp Figure S3. There was differential gene expression between epithelial and mesenchymal cells per treatment condition. (A-C)** Heatmaps display the top 50 differentially expressed genes comparing the E and M clusters for each experimental conditions of **(A)** 21% O_2_, **(B)** 21% O_2_ with 10 ng/mL TGFβ and 50 ng/mL HGF, and **(C)** 1% O_2_ in relation to the clusters from Figure 3A-C.

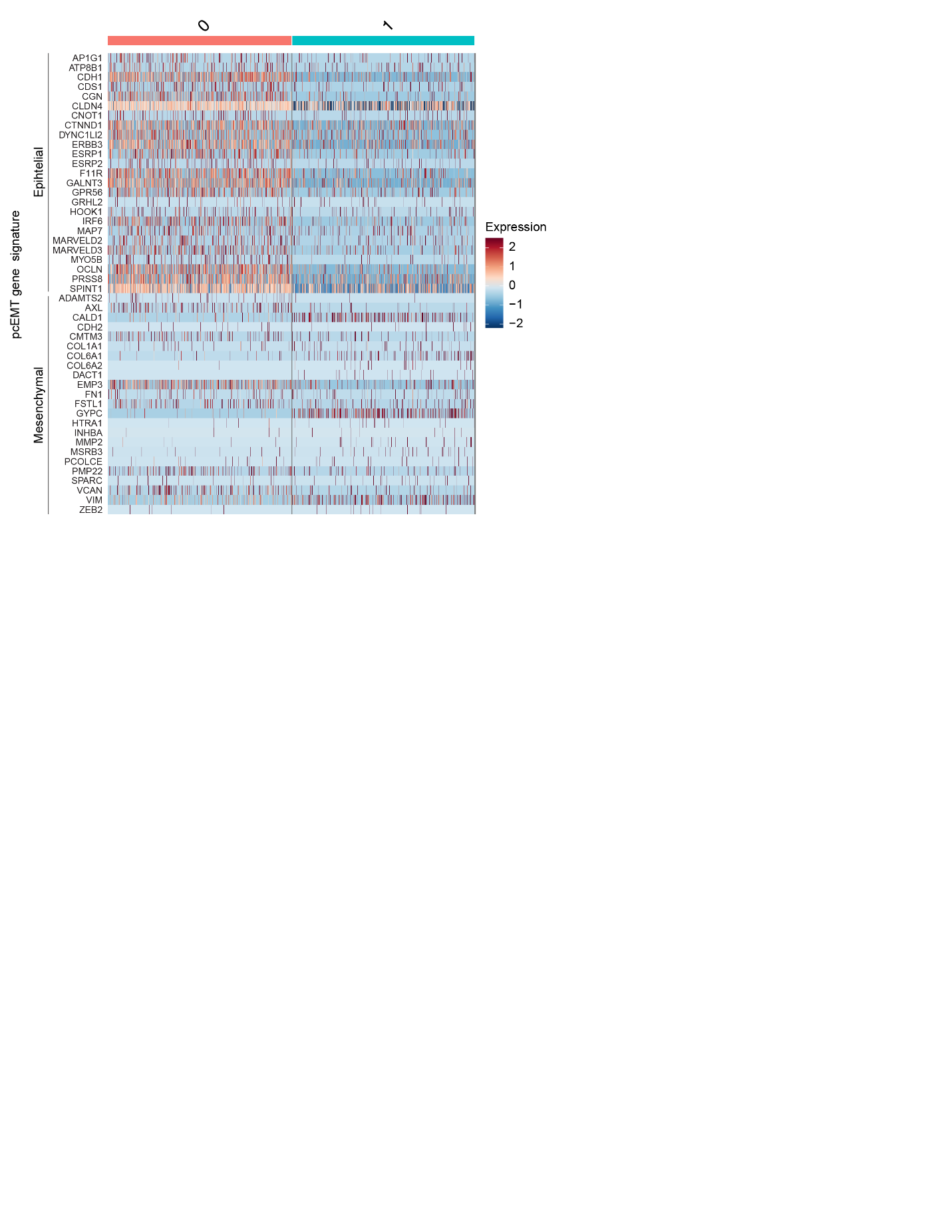

**Supp Figure S4. Patient ductal cells were clustered on EMT gene expression.** The heatmap displays the gene expression of the pcEMT gene signature for the patient ductal cells as clustered in Figure 4C.

**Supp Table S1. Software and algorithms**

| **RESOURCE** | **SOURCE** |
| --- | --- |
| R v4.2.0 | R Development Core Team |
| Bioconductor | (67) |
| cluster v2.1.4 | (68) |
| clusterProfiler v4.4.4 | (69,70) |
| dittoSeq v1.8.1 | (71) |
| dplyr v1.0.9 | (72) |
| escape v1.6.0 | (73) |
| ggplot2 v3.3.6 | (74) |
| ggstatsplot v0.9.4 | (75) |
| limma v3.52.2 | (76) |
| org.Hs.eg.db v3.15.0 | (77) |
| patchwork v1.1.2 | (78) |
| rlang v1.0.5 | (79) |
| scRNABatchQC v0.10.4 | (62) |
| Seurat v4.1.1 | (80-83) |
| stringr v1.4.1 | (84) |
| UCell v2.0.1 | (64) |

**Supp Table S2. qRT-PCR Primers**

| **Target** |  | **Sequence (5' → 3')** |
| --- | --- | --- |
| *CASC3* | Forward | ACCTCGGAAAGGGCTCTTCTT |
|  | Reverse | CGACCCTCATCCTTCCATAGC |
| *CDH1* | Forward | CATCAGGCCTCCGTTTCTG |
|  | Reverse | GGAGTTGGGAAATGTGAGCA |
| *VIM* | Forward | TCTCTGAGGCTGCCAACCG |
|  | Reverse | CGAAGGTGACGAGCCATTTCC |
| *FAT1* | Forward | TCTCTGAGGCTGCCAACCG |
|  | Reverse | CGAAGGTGACGAGCCATTTCC |
